# Fluoxetine-induced neurogenesis and chronic antidepressant effects requires the dopamine D2 receptor

**DOI:** 10.64898/2026.03.29.715084

**Authors:** Gohar Fakhfouri, Morgane Lemasson, Stella Manta, Quentin Rainer, Mohammad Reza Zirak, Bruno Giros, Jean-Martin Beaulieu

**Affiliations:** Centre Cervo, Université Laval, Québec; Department of Psychiatry, Douglas Mental Health University Institute, McGill University, Montreal, Quebec, Canada; Department of Pharmacology and Toxicology, University of Toronto, Toronto, Ontario, Canada; IUCT Oncopole, Toulouse, France; Food and Drug Laboratories Research Center, Mashhad University of Medical Sciences, Mashhad, Iran; Faculté des Sciences Fondamentales et Biomédicales, Université Paris Cité, INCC UMR 8002, CNRS, F-75006 Paris, France

## Abstract

Major depressive disorder (MDD) is a common psychiatric illness with a high proportion of patients being nonresponsive to therapy. Selective serotonin reuptake inhibitors (SSRI) are widely prescribed for treating depression. Chronic SSRI administration is needed for therapeutic effects, a process implicating in part, increased neurogenesis in the hippocampus. Recent genome wide association studies (GWAS) identified the *DrD2* locus, which encodes the dopamine D2 receptor (D2R) as a major risk factor in MDD. Here we demonstrate that behavioural effects associated with chronic administration of the SSRI drug fluoxetine and its accompanying neurogenic effects require D2R. Administration of fluoxetine to congenital D2R-knockout mice, or co-administration of the antidepressant with the antipsychotic D2R antagonist drug haloperidol prevented the neurogenic effects of fluoxetine. Furthermore, while acute behavioural responses to fluoxetine did not require D2R, this receptor was essential for the behavioural effects of chronic fluoxetine. The neurogenic impact of chronic fluoxetine was further associated with beta-arrestin 2-mediated signalling and the hippocampal regulation of the pro-neurogenic factor BDNF. These results support a role of D2R in regulating the therapeutically relevant chronic effects of fluoxetine on mood, BDNF signalling, and associated hippocampal neurogenesis. Furthermore, our findings suggest an unappreciated interaction between genetic risk for MDD and treatment responsiveness as well as a negative interaction between SSRIs and antipsychotic drugs in the regulation of hippocampal neurogenesis.

## 1. Introduction

Major depressive disorder (MDD) is a debilitating psychiatric condition with a high prevalence and largely unmet therapeutic needs (Faquih et al., 2019; Oliveira-Maia et al., 2024; Uyar and Gonul, 2025). Most current antidepressants, including fluoxetine and other selective serotonin reuptake inhibitor (SSRI), target serotoninergic neurotransmission, acutely increasing serotonin bioavailability within minutes (David et al., 2003). However, SSRIs require several weeks to demonstrate clinical efficacy, with more than 30% of patients being refractory to treatment (Rush et al., 2006; Touloumis, 2021). Chronic administration of SSRIs and other classes of antidepressants has been associated with increased neurogenesis in the adult hippocampal dentate gyrus (Lino de Oliveira et al., 2020; Malberg et al., 2000; Wang et al., 2008), a process proposed to account for some of their anxiolytic/antidepressant delayed activity in experimental animals (Alonso et al., 2024; David et al., 2009; Surget et al., 2008). This is consistent with structural changes in the hippocampus, including atrophy observed in depressed patients (Alper et al., 2023; Sheline et al., 1996; Taylor et al., 2014) and animal models of depression (Banasr et al., 2011).

Accumulating evidence now implicates dysregulation of dopaminergic neurotransmission in the etiology of mood disorders, including depression (Grace, 2016; Grotzinger et al., 2025) and antidepressant treatment is associated with dopamine D1 receptor upregulation/activation (Kobayashi et al., 2012), which has been shown to contribute to fluoxetine-induced neurogenesis (Shuto et al., 2018). Furthermore, beta-arrestin 2 (βArr2), a multifunctional adaptor protein involved in G-protein coupled receptor (GPCR) trafficking and signalling is essential for behavioural effects of fluoxetine (David et al., 2009). Interestingly, βArr2 mediates lithium action, a mood stabilizer commonly prescribed in bipolar disorder (Beaulieu et al., 2008), and transduces dopamine D2 receptor (D2R) signalling in the brain by modulating the activity of the Akt/Glycogen synthase kinase 3β (Akt/GSK3β) pathway (Beaulieu et al., 2005). Genome-wide association studies (GWAS) have repeatedly shown that genetic variants of the *DrD2* locus, which encodes D2R, are associated with major depression (Howard et al., 2019; Wray et al., 2018). When positional mapping approaches were used, the D2R gene displayed the strongest evidence of association with depression out of 1,568 genes identified. In addition, using the Psychiatric Omnilocus Prioritization Score (PsyOPS) method, D2R was among the genes with the highest prioritization scores (Adams et al., 2025). Given the central role of D2R in neuropsychiatric disorders, including depression, understanding its potential contribution to antidepressant activity is highly relevant but remains largely unexplored.

Here, we investigate whether D2R expression and signalling are required for fluoxetine’s long-term effects on hippocampal neurogenesis and behaviour. We also assess the specificity of this mechanism by comparing fluoxetine to the monoamine oxidase (MAO) inhibitor tranylcypromine and by distinguishing between behavioural tests with different neurogenic dependencies (David et al., 2009; Santarelli et al., 2003). Our findings highlight D2R as a key effector specifically engaged by chronic, but not acute, fluoxetine treatment *in vivo*. They also suggest that this D2R-mediated effect involves βArr2 signalling and the regulation of hippocampal pro-neurogenic Brain Derived Neurotrophic Factor (BDNF), thereby linking dopaminergic neurotransmission to adult neurogenesis.

## 2. Experimental procedures

### 2.1. Animals

Animal care and handling was performed according to the Canadian Council on Animal Care guidelines and approved by the Animal Care Committees of Universite Laval and the Douglas Research Centre. Animals were housed in groups of 4–5 animals per cage and maintained under standard laboratory conditions: 22° ± 1°C, 60% relative humidity and a regular 12-hour light-dark cycle (7:00–19:00 light on period) with free access to food and water. Mice used throughout the study were from both sexes aged 3-4 months. Groups were age- and sex-matched. Fluoxetine (a serotonin-selective reuptake inhibitor, 18mg/kg/day) was given in the drinking water for 28 days, tranylcypromine (a monoamine oxidase inhibitor, 10mg/kg/day) was injected intraperitoneally for 28 days, and haloperidol decanoate, a long-acting form of haloperidol (a D2R antagonist, 20 mg/kg), was injected once intramuscularly (Röpke et al., 2014). All animals used were on a congenic C57Bl/6J background.

### 2.2. Forced swim test

Mice were dropped into an acrylic glass cylinder (height 25 cm, diameter 9 cm) filled with water at a temperature of 21–23°C. Behavioural despair was measured using immobility time during a 6-min test. Because little immobility occurs during the first 2 min of the test, we recorded immobility only during the remaining 4 min as well as swimming and climbing behaviours. Immobility was scored only when the mice ceased struggling and remained floating and motionless, making only the movements necessary to keep their heads above water.

### 2.3. Novelty suppressed feeding

The test was conducted in an open field (45 × 45 × 45 cm) with a sawdust-covered floor under white illumination (40 W, approximately 2400 lux) positioned immediately above the centre of the open field. The mice were food deprived for 24 hours before testing. At the start of the test, we placed a single food pellet on a round piece of white paper (12.5 cm diameter) at the centre of the apparatus. Each mouse was placed in a corner of the open field with its head directed toward the wall, and the latency to eat was recorded up to a maximum testing period of 10 min (David et al., 2009). Immediately afterwards, we transferred each animal to its home cage for 3 min, and we measured the amount of food consumed to assess changes in appetite as a confounding factor.

### 2.4. BrdU Immunochemistry

Neurogenesis was assessed using the thymidine analog bromodeoxyuridine (BrdU) as a marker for dividing cells (Malberg et al., 2000). Mice were administered with BrdU (75 mg/kg, i.p. dissolved in saline) every two hours (4 times total) and sacrificed 2h after the last BrdU injection. Coronal brain sections (40 um) were obtained from the entire rostrocaudal extension of the hippocampus using a vibratome and stored in PBS with 0.1% NaN3. Every sixth free floating section was rinsed, blocked and probed overnight with anti-mouse BrdU (MAB 3222, 1:500). The labeling was visualized with DAB. An experimenter blinded to the genotype and treatment counted BrdU^+^ cells or cell clusters in the subgranular zone (SGZ) of the dentate gyrus in 12 sections from each mouse. The total count per mouse was obtained by multiplying the sum of counts in all 12 sections by 6.

### 2.5. Immunoblotting

Western blot analyses were performed as previously described (Beaulieu et al., 2008; Beaulieu et al., 2004). Briefly, mice were euthanized by decapitation and the heads were immediately cooled down by immersion in liquid nitrogen for 3 s. The hippocampi were rapidly dissected out on an ice-cold metal surface, snap-frozen in liquid nitrogen and kept at -80 °C until further analysed. Tissues were homogenized in boiling 1% SDS solution for 5 min and total protein concentrations were measured using a DC-protein assay (Bio-Rad). Hippocampal proteins (30 ug) were diluted in Laemmli buffer, separated on 10% Tris-glycine minigels (Invitrogen) and transferred onto nitrocellulose. Membranes were cut according to the molecular weights of the bands of interest, blocked in 5% skim milk for 1h, and probed with primary antibodies (mouse anti-Akt (pan), 2920, CST; rabbit anti-phospho-Akt (T308), 9275, CST; mouse anti-GSK3α/β, SC-7291, SantaCruz; rabbit anti-phospho-GSK3α/β, 9331S; mouse anti-β-actin, Millipore) at 4 °C overnight and revealed using appropriate 680 or 800 nm IR dye-labeled secondary antibodies (Licor Biotechnology, Lincoln, NE). Quantification of bands was carried out by measuring IR dye fluorescence signal using an Odyssey Imager (Licor Biotechnology). Respective total protein signals were used as loading controls for each phospho-protein and treatment-induced changes in phosphorylation of Akt and GSK3β are presented as a fraction of phospho/total ratios in vehicle treated mice. Total Akt and GSK3β signals were also normalized to corresponding β-actin bands as loading controls in WT (n=5 per group).

### 2.6. Bioluminescence resonance energy transfer

Bioluminescence resonance energy transfer (BRET) was employed to monitor beta-arrestin 2 (βArr2) recruitment to D2R in HEK293T cells, maintained in DMEM (Gibco) supplemented with 10% fetal bovine serum at 37 °C with 5% CO2. Expression plasmids encoding human D2R (long isoform, D2L) fused to Renilla luciferase 8 (Rluc8) at the C-terminus (D2L-Rluc8) and human βArr2 fused to mVenus at the N-terminus (venus-βArr2) used in the assay were described elsewhere (Clayton et al., 2014; Guo et al., 2003). HEK293T cells were grown to 80% confluence and 2 million cells were seeded onto 10 cm cell culture dishes. After 24h, transient transfection was carried out using a polyethylenimine (PEI; linear, Mw 25 kDa; PolySciences Inc.)/DNA ratio of 3/1 and the following amounts of plasmids: D2L-Rluc8 (0.2 µg), venus-βArr2 (8 µg) and bovine GRK2 (5 µg). 48h later, cells were washed, resuspended in tyrode’s solution (Sigma), supplemented with 25 mM Hepes (pH 7.4, Sigma) and dispensed in a white 96-well flat bottom plate. Fluoxetine or tranylcypromine (150 µM) or vehicle (tyrode) was added to designated wells and the plate was incubated at 37 °C for 30min. Coelenterazine h (NanoLight Technology, 5 µM) was added and after 5min cells were stimulated with quinpirole (Tocris). BRET1 signal was acquired by calculating the ratio of the light emitted by Venus (535 nm) over that emitted by Rluc8 (485 nm) using a TriStar² Microplate Reader (Berthold). ΔBRET was calculated by subtracting values of non-stimulated cells from those of stimulated conditions. Dose-response curves are presented as percent of βArr2 recruitment in response to the highest dose of quinpirole used and illustrate the average of 3 independent experiments.

### 2.7. ELISA

Hippocampal BDNF content was measured using ELISA. Briefly, a 96-well plate was coated overnight with mouse anti-BDNF monoclonal antibody (GF35L, Calbiochem, CA) at 1ug/ml at 4C. After 3 washes, wells were blocked with 3% BSA/PBS for 2:30 h. 100 µL of hippocampal lysates with determined protein concentrations or standard BDNF solutions were added and shaken for 2h. After 5 washes, samples were probed with 100 µL of polyclonal chicken anti-BDNF antibody (Promega, G164A) at 2.5 µg/ml in 3% BSA/PBS for 2.5h. HRP-conjugated anti-chicken antibody at 1ug/ml in 3% BSA/PBS was added to the wells following 5 washes and samples were further incubated for 1h. After 5 washes, 100 µL of the HRP substrate tetramethylbenzidine (TMB) was added for 15min and the reaction was stopped by adding 100 µL of 1N HCL. The absorbance was read at 450 nm within 30 min of HCL addition. From the time of sample loading, all the steps were performed at RT on a shaker. The washing steps were done with PBS supplemented with 5% Tween-20 (PBS-T). BDNF concentrations were calculated from the standard curve and expressed as pg/µg of total protein.

### 2.8. Data analysis and statistics

Results are presented are means ± SEM. All analyses were carried out using GraphPad Prism 7 or 10. The significance of differences between two groups was determined by unpaired student’s *t* test (two-tailed). For multiple comparisons, one-way or two-way ANOVA followed by Tukey’s post-hoc was employed, unless otherwise specified. Dose-response data were analyzed using nonlinear regression by fitting data to a sigmoidal dose-response equation. *p* <0.05 was considered significant.

## 3. Results

### 3.1. Dopamine D2 receptor is necessary for pro-neurogenic effects of chronic fluoxetine

We explored whether the neurogenic action of chronic fluoxetine could be contingent on D2R. Fluoxetine treatment for 28 days induced proliferation of the hippocampal progenitor cells in WT mice, as evidenced by a dramatic increase in the number of BrdU positive cell clusters in the adult subgranular zone (SGZ) (*p*=0.0004) (Fig. 1A; 1S). Subzonal analysis of the SGZ (Fig. 1B) revealed that the neurogenic effect was exerted in both ventral and dorsal regions (*p*=0.0074 and *p*=0.0075 vs vehicle, respectively). Chronic fluoxetine failed to affect neurogenesis in the dentate gyrus of D2KO mice (*p*=0.7338) (Fig. 1A). D2R ablation, *per se*, led to an elevated number of BrdU positive cell clusters in the SGZ (*p*=0.0007 vs vehicle-treated WT). The impact of D2R knockout on neurogenesis was robust in the dorsal and ventral zones (*p*=0.0061 and *p*=0.0278 vs vehicle, respectively) (Fig. 1B).

**Fig. 1.**
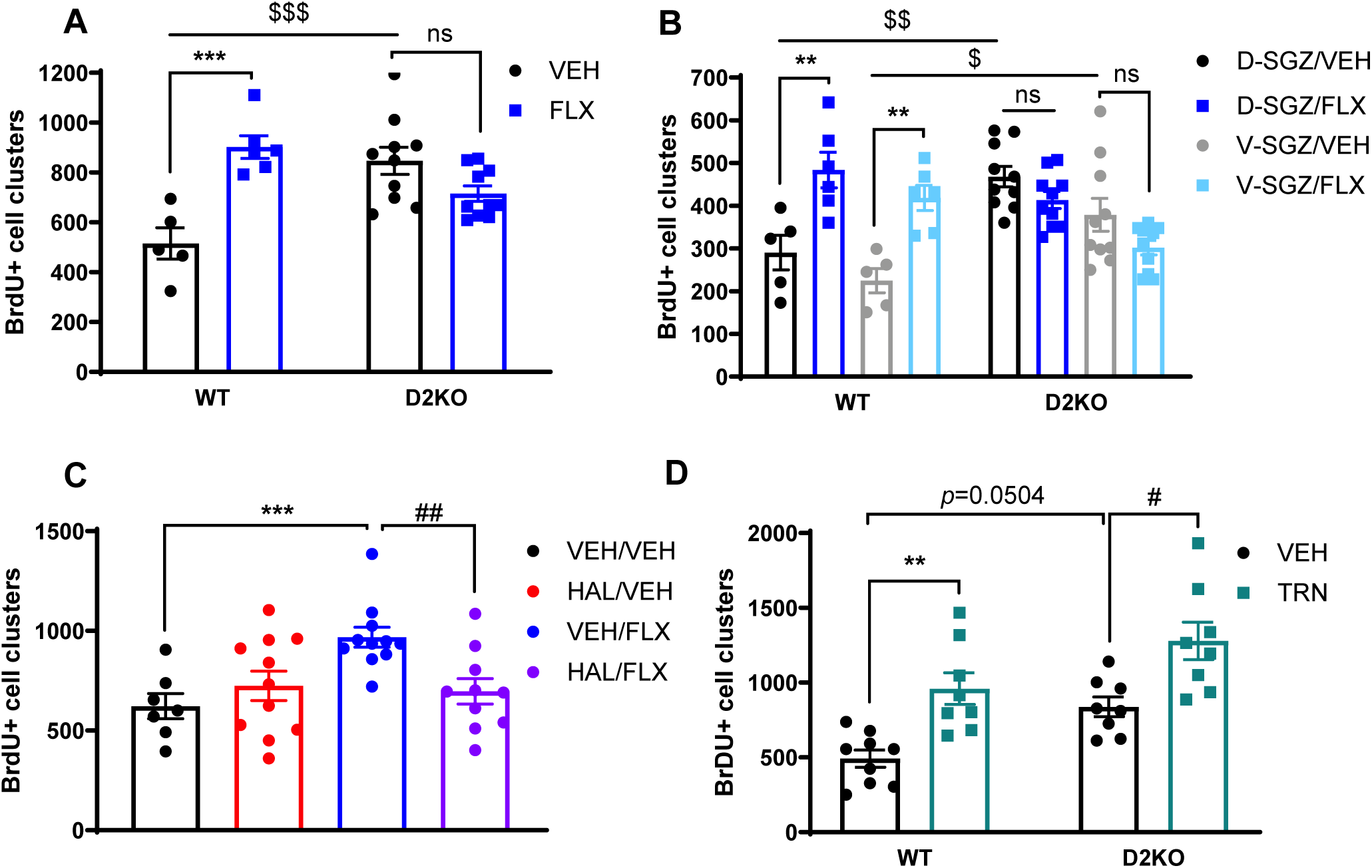
D2R is essential for fluoxetine-induced hippocampal neurogenesis. WT or dopamine D2 receptor knockout (D2KO) mice were treated for 28 days with different regimens. On day 28, BrdU (75 mg/kg) was injected intraperitoneally every 2h for 4 times. Mice were perfused 2h after the last BrdU injection to examine the effects of the chronic treatment on dentate gyrus neurogenesis. **(A)** Chronic FLX increased the number of BrDU positive cell clusters in the SGZ (n=5-6), **(B)** in both ventral and dorsal regions. D2R deletion led to an elevated number of BrDU positive cell clusters (n=10) **(A)**, with the impact being robust in both dorsal and ventral zones **(B)**. FLX, however, failed to induce proliferation of the hippocampal progenitor cells in D2KO mice (n=10) **(B)**. **(C)** A sustained release form of haloperidol (HAL, 20mg/kg single intramuscular injection) in WT mice abolished fluoxetine-induced SGZ neurogenesis (n=10). HAL alone did not affect the BrdU labeling in the dentate Gyrus (n=7-11). **(D)** Tranylcypromine (TRN, 10 mg/kg/day for 28 days) increased BrDU+ cell clusters in the SGZ of both WT and D2KO mice (n=8-9). Data are expressed as the mean ± SEM of the BrdU-positive cell cell clusters for the SGZ. \*\**p*<0.01, \*\*\**p*<0.001 versus WT treated with VEH; $*p*<0.05, $$*p*<0.01 and $$$*p*<0.001 versus D2KO treated with VEH; ## *p*<0.01 versus WT treated with FLX. # versus D2 KO treated with VEH.

To verify whether the absence of fluoxetine-induced neurogenesis in D2KO mice was due to a developmental adaptation, we directly exerted a chronic blockade of D2R in WT mice using intramuscular delivery of slow-release haloperidol decanoate (20mg/kg). Pharmacological antagonism of D2R by haloperidol abolished fluoxetine-induced SGZ neurogenesis (*p*=0.004), replicating the lack of fluoxetine neurogenic action in D2KO mice and confirming the requirement of functional D2R activation for fluoxetine-induced neurogenesis in the adult hippocampus (Fig. 1C).

Interestingly the impact of D2R activation on hippocampal neurogenesis was more important than that of the direct main pharmacological target of fluoxetine, the serotonin transporter (SERT). Indeed, treatment with fluoxetine had a similar efficacy to stimulate neurogenesis between SERT-KO and their WT littermates (number of BrdU positive cells in SERTKO: 1.57 fold basal, *p*=0.0212 versus vehicle; in WT 1.48 fold basal, *p*=0.0223 vs vehicle) (Fig. 2S). There was no difference in genotype (main effect F(1, 32)=1.698; *p*=0.2019) and basal levels of BrdU positive cells in the SGZ were similar between the two genotypes (*p*=0.7683). This observation is in line with previous reports indicating that the SERTKO genotype, *per se*, does not affect the proliferation of adult stem cells or survival of newborn cells (Benninghoff et al., 2012; Schmitt et al., 2007).

**Fig. 2.**
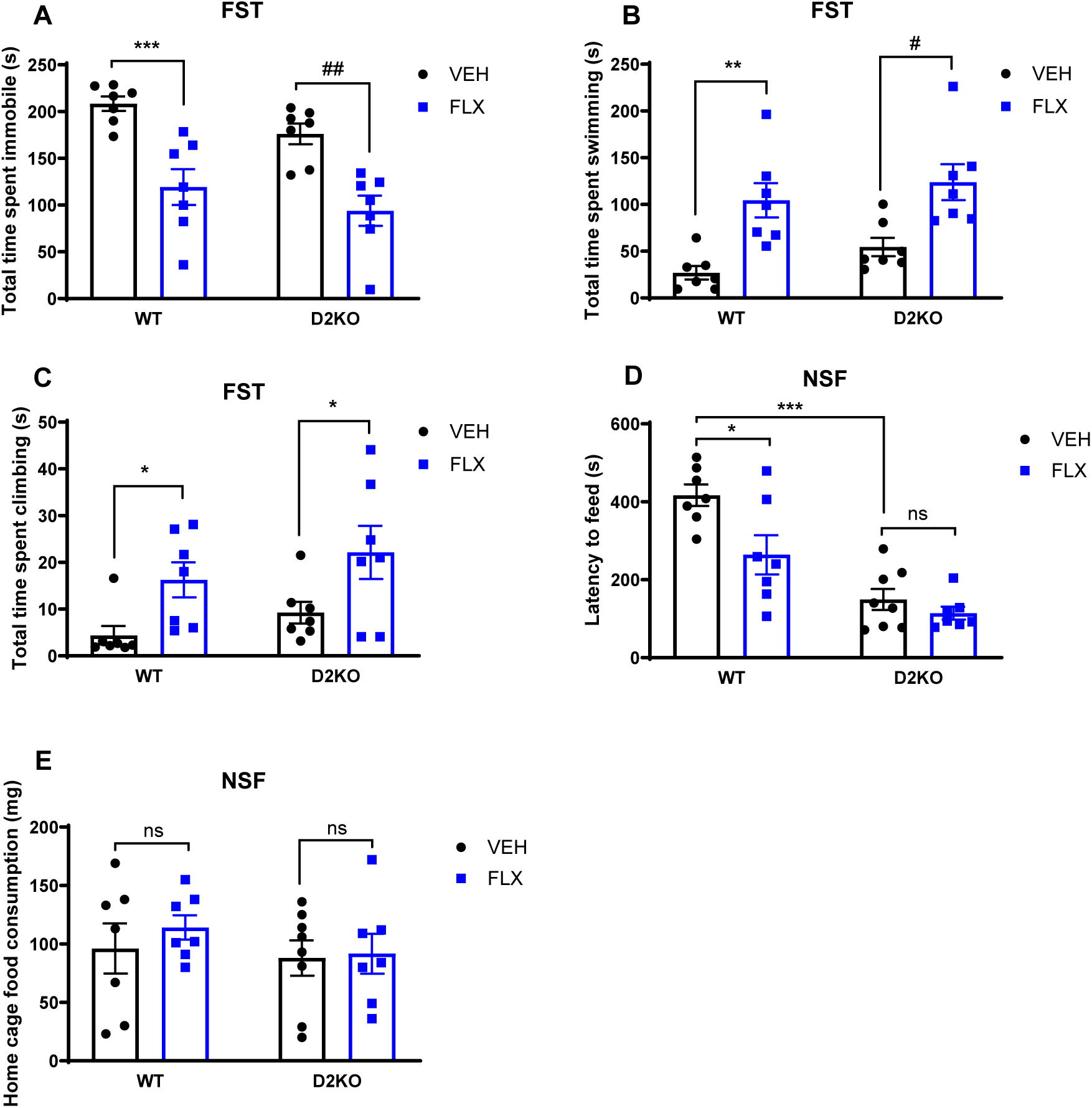
D2R genotype and fluoxetine differentially affect performance in forced swim test and novelty suppressed feeding. **(A-C)** Forced swim test (FST) in wild type and D2R knockout mice after a single fluoxetine injection. D2R knockout (D2KO) mice and wild type (WT) littermates were injected intraperitoneally with either 0.9% NaCl (VEH) or fluoxetine (FLX, 18 mg/kg dissolved in 0.9% NaCl) and tested in forced swim test 1h after injection. **(A)** Total time spent swimming; **(B)** immobile and **(C)** climbing was measured. D2R genotype does not alter performance in the FST. Fluoxetine significantly reduced immobility and enhanced swimming and climbing in both WT and D2KO mice. **(D-E)** Novelty suppressed feeding (NSF) test in WT and D2KO mice after chronic fluoxetine administration. WT and D2KO mice received either standard drinking water (VEH) or fluoxetine (FLX, 18 mg/kg/day) in drinking water during 28 days and tested in the NSF (n=7 per group). **(D)** Latency to eat was measured after 24 hours of food deprivation. In WT mice, fluoxetine significantly shortened the food consumption latency, but had no effect in D2KO mice. D2KO genotype resulted in a drastic reduction of latency to feed. **(E)** After the test, mice were allowed to return to their home cage where food consumption was measured for 3 minutes. Consumption in home cage was not affected by genotype or treatment. Values are expressed as the mean ± SEM. \**p*<0.05, \*\**p*<0.01, \*\*\**p*<0.001 versus WT treated with VEH; #*p*<0.05 and #*p*<0.01 versus D2KO treated with VEH.

Lastly, we administered the monoamine oxidase (MAO) inhibitor tranylcypromine, known to increase neurogenesis in animal models to WT and D2KO mice. In agreement with previous observations (Malberg et al., 2000) chronic tranylcypromine (10mg/kg for 28 days) increased BrdU^+^ cell clusters in the SGZ of WT mice (*p*=0.0005). But unlike fluoxetine, tranylcypromine produced a similar effect on hippocampal neurogenesis in D2KO mice (*p*=0.0109 vs vehicle), suggesting that despite sharing pro-neurogenic effects, the D2R mediation of this effect is unique to fluoxetine and not a common property of antidepressant treatment (Fig. 1D).

### 3.2. Dopamine D2 receptor is not required for antidepressant-like effects of acute Fluoxetine

To assess the requirement of D2R for the acute antidepressant-like effect of fluoxetine, a single dose (18mg/kg, i.p.) was administered to WT and D2KO mice and behaviour was evaluated 1h later in the FST. Acute fluoxetine significantly reduced behavioural despair across all behavioural measures in the FST (immobility, F(1, 24)=36.36, *p*<0.0001; swimming, F(1, 24)=25.35, *p*<0.0001; climbing, F(1, 24)=10.98, *p*=0.0029). In WT genotype, fluoxetine significantly decreased the immobility time (*p*=0.0009) and increased swimming duration (*p*=0.0051). Fluoxetine induced a trend toward increase in climbing time, which was not significant with Tukey’s *post-hoc* test but reached significance with Fisher’s LSD (*p*=0.034). Importantly, no genotype effect was observed and in D2KO mice fluoxetine exhibited a significant anti-depressive-like effect similar to that in WT littermates, namely reducing immobility (*p*=0.0022) while prolonging swimming (*p*=0.0128) and climbing (*p*=0.0225, Fisher’s LSD) (Fig. 2A-C). These results indicate that the acute antidepressant-like effects of fluoxetine in mice occur independently of D2R receptor.

### 3.3. Dopamine D2 receptor is required for antidepressant effects of chronic Fluoxetine in the novelty-suppressed feeding test

The novelty suppress feeding test (NSF) test possesses the relevant property of responding to chronic, but not acute, antidepressant treatment (Belovicova et al., 2017). Chronic fluoxetine (18 mg/kg/day for 28 days) had a significant effect in latency to feed in the novel environment (main effect, F (1, 25)=8.285, *p*=0.0081). Chronic fluoxetine significantly decreased the latency to consume food in WT (*p*=0.0157). Corroborating the neurogenesis observations, however, the same treatment failed to affect the latency in D2KO littermates (*p*=0.8691). Two-way ANOVA revealed a significant effect of D2R ablation on this measure (F (1, 25)= 41.01, *p*<0.0001) (Fig. 2D). Interestingly, untreated D2KO mice exhibited a significantly shorter feeding latency relative to untreated WT littermates (*p*<0.0001), indicating an anti-depressive/anxiolytic phenotype in D2KO mice (Fig. 2D), conforming to the pro-neurogenic effect associated with this genotype (Fig. 1B-C). There was, however, no treatment by genotype interaction (F (1, 25) = 3.288, *p*=0.0818). The home cage food consumption was unaltered regardless of genotype and treatment (Fig. 2E). These findings demonstrate the requirement of D2R for behavioural effects of neurogenetic-dependent chronic fluoxetine in NSF and the development of anti-depressive phenotype in D2R absence.

### 3.4. Chronic fluoxetine regulates pro-neurogenic βArr2-mediated Akt/GSK3β signalling in the hippocampus

Activation of D2R regulates the Akt/GSK3β (Beaulieu et al., 2005), which is also involved in neuronal progenitor cell survival (Del Puerto et al., 2023) and in the signalling of the pro-neurogenic factor BDNF (Atwal et al., 2000; Jun et al., 2012; Sairanen et al., 2005). This pathway is also targeted by antidepressants, lithium and other mood stabilizers (Beaulieu et al., 2009; Beaulieu et al., 2008; Del’ Guidice and Beaulieu, 2015) thus providing a potential nexus for the interaction of fluoxetine, adult neuronal proliferation and D2R signalling. We first investigated the impact of chronic fluoxetine treatment on Akt/GSK3β signalling in the hippocampus of WT C57Bl/6J mice that were administered fluoxetine or vehicle for 28 days (Fig. 3S). Western blot analysis of hippocampal tissue from WT mice showed, no effect of treatment on the total Akt and GSK3β expression (*p*=0.4821 and *p*=0.7387 vs vehicle (Fig. 3A). In contrast, chronic fluoxetine induced an increase phosphorylation of GSK3β at S9 (*p*=0.0025 vs vehicle) and a similar increase in phosphorylation of its regulating kinase Akt at T308 (*p*=0.0265 vs vehicle) in response to fluoxetine (Fig. 3A).

**Fig. 3.**
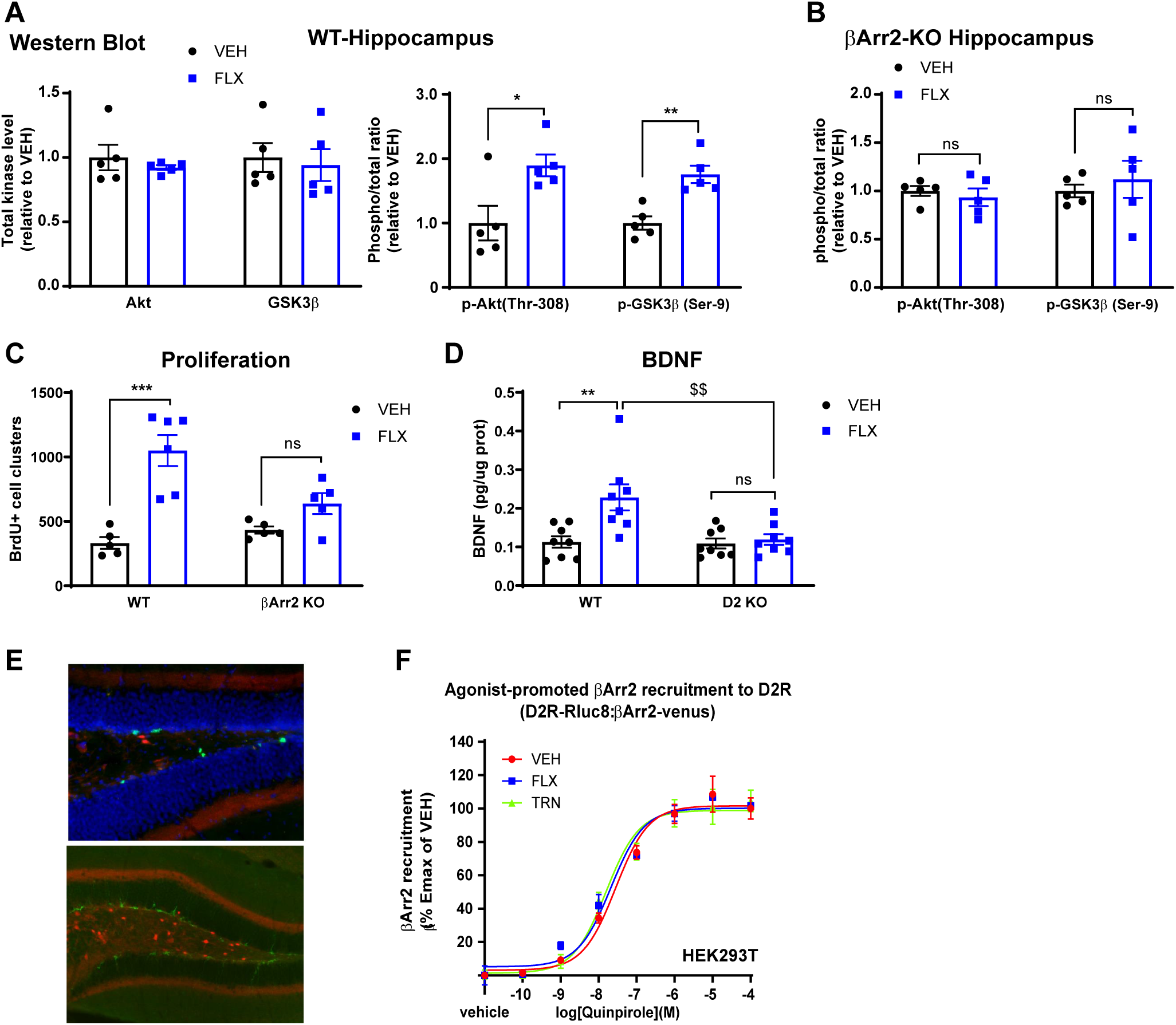
Fluoxetine relies on βArr2-mediated D2R signalling to induce hippocampal effects. Mice were treated for 28 days with either 0.9% NaCl (VEH) or fluoxetine (FLX, 18 mg/kg in 0.9% NaCl). **(A)** Chronic FLX did not change total Akt and GSK3β expression (left) but increases phosphorylation of Akt at Thr308 and its substrate GSK3β at Ser9 in WT mouse striatum (right). **(B)** The effect of FLX on Akt at Thr308 and GSK3β at Ser9 was loss in βArr2KO mice. **(C)** On day 28, BrdU (75 mg/kg) was injected intraperitoneally every 2h for 4 times. Mice were perfused 2h after the last BrdU injection to examine the effects of the chronic FLX treatment on dentate gyrus neurogenesis. **(D)** FLX significantly increases hippocampal contents of BDNF in WT mice, but not in D2KO littermates **(E)** Immunostaining of the dentate gyrus from D2tomato mice for specific markers (in green) shows D2R expression is not detectable on neural progenitor cells (Ki-67 staining, upper panel) or newly born neurons (DCX staining, lower panel) of the dentate gyrus, micrographs are at 20x magnification **(F)** FLX and tranylcypromine (TRN) do not affect quinpirole-mediated βArr2 recruitment to D2R as evaluated using BRET sensors transfected in HEK293T cells. \**p*<0.05, \*\**p*<0.01, \*\*\**p*<0.001 versus WT treated with VEH; $$*p*<0.01 versus D2KO treated with FLX.

Chronic fluoxetine alters βArr2 gene expression in the hippocampus in a mouse model of anxiety/depressive-like states (David et al., 2009). Consistently, genetic ablation of βArr2 was reported to abolish the antidepressant impact of chronic fluoxetine in the NSF (David et al., 2009), which relies on the presence of intact progenitor cells in the hippocampal SGZ (Santarelli et al., 2003). Investigation of the impact of chronic (28 days) fluoxetine administration on Akt/GSK3β signalling in the hippocampus of mice lacking βArr2 showed no effect of fluoxetine in βArr2KO mice (*p*=0.9251 and *p*=0.3322 vs vehicle, respectively for pGSK3β and pAkt ratios) pointing to the requirement of βArr2 for fluoxetine-driven Akt/GSK3β modulation (Fig. 3B, Fig. 3S). We than assessed neurogenesis in this region in response to chronic fluoxetine in βArr2KO mice. As opposed to its pronounced neurogenic properties in WT mice, long-term fluoxetine administration did not alter the number of BrdU^+^ cell clusters in the SGZ of βArr2 deficient mice (*p*=0.3562 vs vehicle, Fig. 3C), mirroring the lack of behavioural response in NSF (David et al., 2009) and further implicates βArr2 in chronic fluoxetine effects.

### 3.5. D2R mediates the upregulation of hippocampal BDNF by chronic fluoxetine

Brain-derived neurotrophic factor (BDNF), has been implicated in various features of adult hippocampal neurogenesis including proliferation (Jun et al., 2012) and long-term survival of newborn hippocampal neurons (Sairanen et al., 2005). Chronic, but not acute, antidepressant treatment from different classes upregulate BDNF (Nibuya et al., 1995), suggesting it as a potential mediator of antidepressant action (Jun et al., 2012). To investigate whether D2R contributes to fluoxetine-induced neurogenesis through BDNF, we measured the hippocampal BDNF level after chronic administration of fluoxetine (Fig. 3D). ELISA revealed that chronic administration of fluoxetine was associated with elevated BDNF concentrations in the WT hippocampus (*p*=0.0026 vs vehicle, n=8). In contrast, fluoxetine failed to modulate BDNF levels in D2RKO mice (*p*=0.9836 vs vehicle, Fig. 3D, n=8), mirroring lack of an antidepressant effect in behaviour and neurogenic response at the cellular level in D2RKO mice. This result suggests that D2R-mediated BDNF upregulation in the hippocampus contributes to the D2R-dependent behavioural and neurogenic dimensions of chronic fluoxetine.

### 3.6. Fluoxetine does not regulate D2R signalling directly in neuronal progenitor cells

To investigate the possibility that fluoxetine may engage D2R signalling in neuronal progenitor cells of the SGZ, we first used reporter mice expressing tomato D2R expressing cells. Labeling of sections of the dentate gyrus obtained from these mice to identify either progenitor cells (Ki-67 positive) or newly born neurons (Doublecortin positive) indicated that D2R expression is restricted to interneurons in the dentate gyrus and is not expressed in progenitor cells (Fig. 3E).

In addition, we evaluated the impact of fluoxetine on βArr2 recruitment to D2R as a correlate of receptor activation in transfected cells using a BRET sensor system (Clayton et al., 2014; Guo et al., 2003). As it may be expected from binding studies (Cusack et al., 1994; Penttilä et al., 2004), addition of Fluoxetine to the transfected cells had no impact on βArr2 recruitment to D2R (Fig. 4S). Moreover, neither fluoxetine nor tranylcypromine, affected Quinpirole-promoted βArr2 recruitment to D2R (Fig. 3F) in this system, thus demonstrating that fluoxetine does not engage D2R/βArr2 mediated signalling by directly acting on D2R.

## 4. Discussion

Results presented here demonstrate that D2R is essential for the expression of behavioural and neurogenic effects of chronic fluoxetine treatment. Concomitant exposure to the D2R antagonist antipsychotic drug haloperidol or congenital D2R inactivation prevented the neurogenic effects of chronic fluoxetine treatment. Furthermore, chronic fluoxetine had no significant impact on pro-neurogenic hippocampal BDNF levels and neurogenesis-dependent behavioural responses in the NSF test. Importantly, performance in the FST in response to acute fluoxetine was not affected by D2R inactivation, thus indicating a specific contribution of D2R signalling in the therapeutically relevant chronic action of this antidepressant, which is particularly interesting in the context of recent genome-wide association meta-analyses implicating genetic variants of D2R in depression (Howard et al., 2019; Wray et al., 2018).

Chronic fluoxetine in various regimens potentiates neurogenesis in the adult hippocampus in rodents and humans (Micheli et al., 2018) (Boldrini et al., 2009). Increased neurogenesis in experimental animals is a common property of antidepressants from different classes (Malberg et al., 2000). Ablating hippocampal neurogenesis by X-ray irradiation renders antidepressants inactive in behavioural paradigms used to model chronic antidepressant response in mice such as the NSF (David et al., 2009; Dranovsky and Hen, 2006). Moreover, depletion of adult neurogenesis using the chemotherapy drug temozolomide in mice induces behavioural and biological changes relevant to depression (Egeland et al., 2017). The possible role of dopamine neurotransmission in the action of fluoxetine was suggested by the observation that fluoxetine upregulates D1R transcription and signalling as a mechanism responsible for its cellular and behavioural effects (Shuto et al., 2018). Furthermore, fluoxetine, desipramine and tranylcypromine, when given repeatedly increased D2-like but not D1-like receptor sensitivity in the rat, as evidenced by enhanced behavioural response selectively to D2-like agonists (Ainsworth et al., 1998).

Our results underscore a role of D2R in mediating the effects of chronic fluoxetine on hippocampal neurogenesis. This contribution is in addition to the one of the 5HT1A receptor, which was initially reported to be essential for this neurogenic effect (Santarelli et al., 2003). Interestingly, the MAO-A and B inhibitor tranylcypromine retains its pro-neurogenic activity in D2KO mice. This indicates that: 1. the unresponsiveness to fluoxetine does not stem from a ceiling effect of the D2KO genotype; and 2. this D2R-dependent mode of action is not a common property of different classes of antidepressants. A similar discrepancy has been observed in mice lacking the 5HT1A receptor, which are still responding to a tricyclic antidepressant stimulating norepinephrine transmission (Santarelli et al., 2003). It is thus possible that drugs like fluoxetine, affecting principally serotonin may have an additional dependence on dopamine transmission as compared to less selective antidepressant drugs.

The dentate gyrus constitutes the only region within the hippocampus where D2R mRNA is detected (Rocchetti et al., 2015). Using a D2R Cre:ribotag mouse line, Puighermanal et al., confirmed that D2R- expressing cells comprise mainly glutamatergic hilar mossy cells and also a subpopulation of GABAergic hippocampal interneurons (Puighermanal et al., 2015). In single cell RNA sequencing, D2R appears in clusters of mossy cells and interneurons but not in principal neurons of the hippocampus (Saunders et al., 2018). We also show that the *drd2* gene is not active in either progenitor cells or newly born neurons of the SGZ (Fig. 3E). Furthermore, fluoxetine itself had no functional pharmacological engagement with the D2R (Fig. 3F and S3), thus pointing to an indirect contribution of D2R involving circuit level modulations.

In addition to neurogenesis, D2R was involved in the regulation of BDNF by fluoxetine (Fig. 3D). Ample evidence implicates BDNF in regulating adult neurogenesis in the SGZ of the dentate gyrus (Lee et al., 2002; Scharfman et al., 2005; Waterhouse et al., 2012) and deletion of its receptor TrkB in adult progenitors increases depression/anxiety-like behaviour (Bergami et al., 2008). BDNF is also involved in long-term synaptic plasticity throughout the brain (Edelmann et al., 2014). BDNF is upregulated by chronic antidepressant treatment (Kozisek et al., 2008) and TrkB receptor in neural progenitor cells governs behavioural sensitivity to antidepressants (Li et al., 2008). In humans, the expression of BDNF and the TrkB receptor is diminished in the hippocampus and PFC of suicide subjects (Dwivedi et al., 2003) and elevated in patients treated with antidepressants (Chen et al., 2001). Our investigation demonstrates that, fluoxetine relies on D2R to increase the hippocampal BDNF content since D2R deletion renders mice insensitive to fluoxetine-induced BDNF upregulation. This is in keeping with the neurogenic and behavioural effects of fluoxetine.

However, BDNF, is not produced by interneurons of the hippocampus (Marty et al., 1996), which are also expressing D2R. Indeed, the pro-BDNF gene is mainly being transcribed in principal neurons and neural precursor cells (Saunders et al., 2018). To reconcile the D2R-dependent BDNF upregulation by fluoxetine with the hippocampal D2R expression pattern, we postulate that modulation of interneuron activity as a consequence of D2R inhibition by fluoxetine regulates target granule neurons in the dentate gyrus to increase BDNF production and release, thereby potentiating the process of neurogenesis in adjacent neural precursor cells in the SGZ. In support of this model, GABAergic interneurons regulate BDNF expression and release from granule cells of the dendate gyrus (Marty et al., 1996). Alternatively, activation of mossy cells by fluoxetine-mediated D2R inhibition likely leads to BDNF release from mossy cell axons into the molecular layer (Hashimotodani et al., 2017), which is sensed by TrkB receptors on neural progenitors cells to promote neurogenesis. Future studies should delineate the nature of the D2R expressing neurons responsible for the observed effects.

The D2R exert its action via parallel Gαi/o G-protein and βArr-mediated mechanisms (Beaulieu and Gainetdinov, 2011). Interestingly, David et al. showed that following long-term corticosterone exposure, which models depression in mice, βArr1 and βArr2 gene transcriptions change differentially across limbic structures and that fluoxetine treatment restores these alterations selectively in the hypothalamus (David et al., 2009). In addition, the neurogenic effects of chronic fluoxetine treatment are lost in βArr1KO mice (Mendez-David et al., 2023). Congenital deletion of the βArr2 gene results in increased consumption latency in drug-naïve mice indicative of a depressive phenotype, and leads to unresponsiveness to behavioural effects of chronic fluoxetine (David et al., 2009). Single nucleotide polymorphisms in the βArr2 gene have been also associated with a lower response to antidepressant treatment in patients (Petit et al., 2018). This agrees with our observation of a lack of pro-neurogenic response to chronic fluoxetine in the SGZ of βArr2KO mice (Fig. 3C), suggesting the requirement of βArr2 for the activity of chronic fluoxetine.

Interestingly, βarr2 is pivotal to the signalling complex that mediates the responsiveness to mood stabilizer lithium (Beaulieu et al., 2008). The complex comprises the serine/threonine kinase Akt, βArr2 and protein phosphatase A2 (PP2A) (Beaulieu et al., 2007a; Beaulieu et al., 2005), and its formation is governed by D2R in the mouse brain (Beaulieu et al., 2004; Beaulieu et al., 2007b). Activation of D2R in dopaminoceptive brain regions promotes the formation of a signalling complex comprised of Akt, βArr2 and protein phosphatase A2 (PP2A), leading to dephosphorylation and inhibition of Akt, thereby dephosphorylation and activation of its substrate GSK3β, which gives rise to DA-related effects (Beaulieu et al., 2005). This pathway is also targeted and inhibited by other mood stabilizers in a βArr2-and D2R-dependent fashion (Del’Guidice et al., 2015), pointing towards the importance of this pathway in mood regulation.

Constitutive knockout of D2R leads to shortening of the feeding latency in the NSF and increased BrdU labeling in the SZG. The dentate gyrus is also bigger in D2KO mice which could be a reflection of higher neurogenesis in the hippocampus (Guma et al., 2018). These features are reminiscent of an antidepressant-like phenotype in D2R deficient mice. Intriguingly, D2R deletion has no impact on the expression of behavioural despair in the FST. This signifies a selective and previously unappreciated role for D2R in the pathways associated with performance in the neurogenesis-dependent NSF test but not the FST.

The observation that fluoxetine relies on βArr2 to exert pro-neurogenic and chronic antidepressant-like actions prompted us to interrogate the impact of fluoxetine on βArr2-mediated Akt/Gsk3β phosphorylation in the hippocampus. Chronic treatment of WT mice with fluoxetine was associated with an increase in the basal phosphorylation of Akt and its effector GSK3β, reminiscent of D2R inhibition. The selectivity of fluoxetine’s effect on this pathway (as opposed to PI3K/Akt downstream of TrkB) was confirmed by its absence in the βArr2KO mice.

The observation that D2R blockade by haloperidol alone did not significantly affect BrdU labeling in the dentate gyrus, as reported by other groups (Halim et al., 2004; Malberg et al., 2000) suggests that fluoxetine’s inhibitory activity on D2R is required but not sufficient for the expression of its chronic properties. It is conceivable that the modulation of different signalling pathways, such as those mediated by SERT, D1R and D2R act in concert to bring about the effects. However, inhibition of SERT alone does not seem to be sufficient, since ablation of SERT does not abolish the neurogenic action of fluoxetine. It is also possible that haloperidol possesses a low potency in modulating the D2R-mediated pathways with a neurogenic outcome. Another possibility is that, in order for fluoxetine to modulate the βArr2-mediated D2R signalling, the orthosteric binding site of D2R should not be occupied by an antagonist and therefore the receptor should not be pre-stabilized in the inactive conformation. The acute behavioural impact of fluoxetine in the FST occurs, however, regardless of the D2R genotype, which concurs with the dispensable role of D2R in this test and further confirms D2R as an indirect target of chronic, not acute, fluoxetine action.

In summary, our study uncovers a previously unrecognized role of D2R in mediating the chronic, but not acute, antidepressant effects of fluoxetine. It also suggests that this D2R-dependent effect involves βArr2 signalling modality and the regulation of hippocampal BDNF, thereby linking dopaminergic neurotransmission to adult neurogenesis. Our findings may have therapeutic potential by providing insight into a possible mechanism that underlies resistance to SSRI treatment in certain patients, where variants in the D2R gene could diminish the therapeutic efficacy of fluoxetine and potentially other SSRIs. Furthermore, the negative interaction between haloperidol and fluoxetine also points toward drug interactions that could have a negative impact on responsiveness to antidepressant drug therapy. This could have important clinical implications considering the prevalent use of antipsychotics in patients with MDD (Gerhard et al., 2018; Zhou et al., 2022). Our results are also consistent with GWAS demonstrating a strong association between the D2R gene and major depression. Collectively, our findings provide novel insights that could contribute to the development of enhanced therapies for mood disorders.

## CRediT authorship contribution statement

All authors conceived experiments and interpreted findings

GF, BG & JMB wrote the manuscript

GF, SM and ML conducted experiments analysed data equally.

## Acknowledgements

Camille Latapy and Annie Barbeau for mouse colony maintenance.

## Supplementary materials

Figures 1-4S

