## Supplemental Figures S1-S4 for "Fluoxetine-induced neurogenesis and chronic antidepressant effects requires the dopamine D2 receptor"

**Fig. 1S.** **D2R is essential for fluoxetine-induced hippocampal neurogenesis**

**Fig. 2S.** **SERT is not necessary for fluoxetine-induced hippocampal neurogenesis**

**Fig. 3S.** **Uncropped western blot membranes of the proteins of interest**

**Fig. 4S.** **Fluoxetine does not act as an agonist at D2R**

**Fig. 1S. D2R is essential for fluoxetine-induced hippocampal neurogenesis**
(A-D) Representative photomicrographs of proliferating BrdU-labeled cells, which were localized to the subgranular zone (SGZ) of the hippocampus.

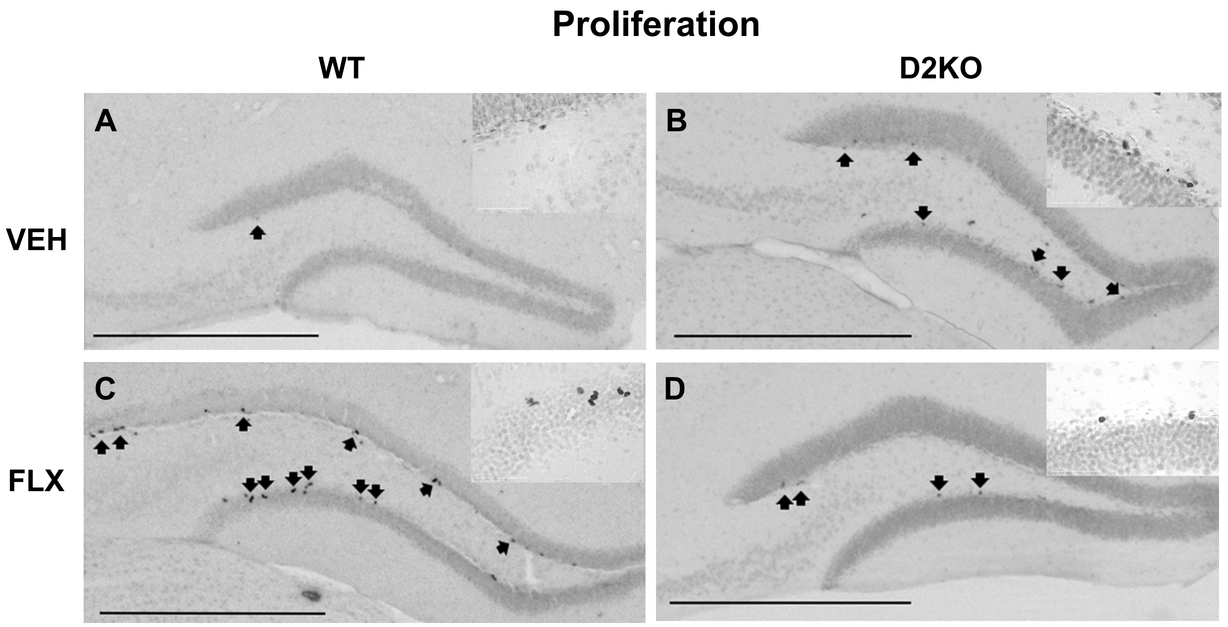


**Fig. 2S. SERT is not necessary for fluoxetine-induced hippocampal neurogenesis**

28 days of Fluoxetine treatment (FLX, 18 mg/kg in 0.9% NaCl) retained its pro-neurogenic action in Serotonin transporter knockout (SERT KO) mice. SERT deletion did not alter basal BrdU labeling in the SGZ as compared to WT (n=8-10). Data are expressed as the mean ± SEM of the BrdU-positive cell counts for the SGZ. * $p<0.05$ versus WT treated with VEH; & $p<0.05$ versus SERT KO treated with VEH.

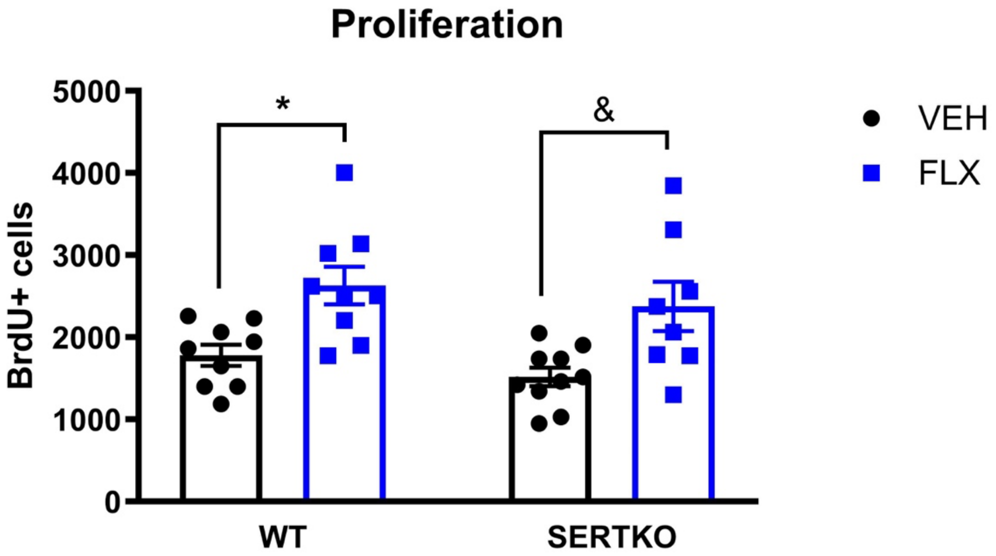

**Fig. 3S. Uncropped western blot membranes of the proteins of interest**

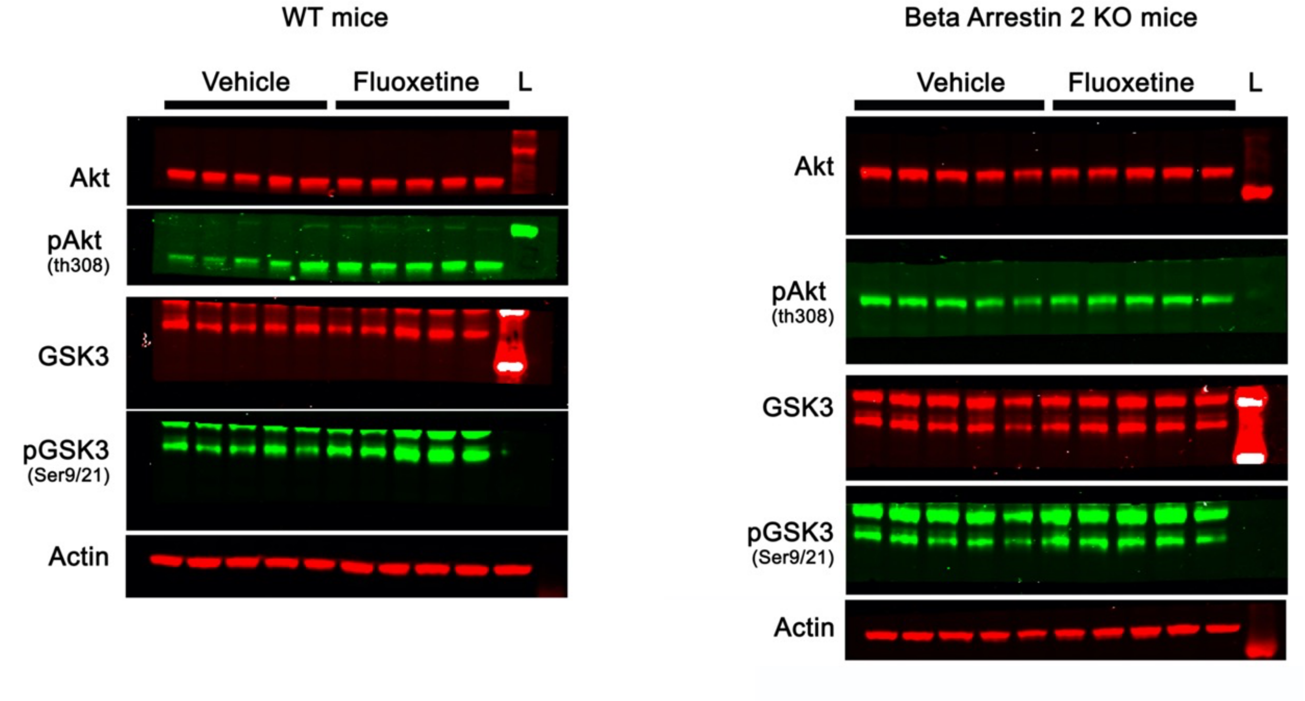


**Fig. 4S. Fluoxetine does not act as an agonist at D2R**

FLX does not induce βArr2 recruitment to D2R as evaluated using BRET sensors transfected in HEK293T cells. Data are presented as fold change in BRET ratio following FLX addition (D2L-Rluc8:venus-βArr2, n=3).

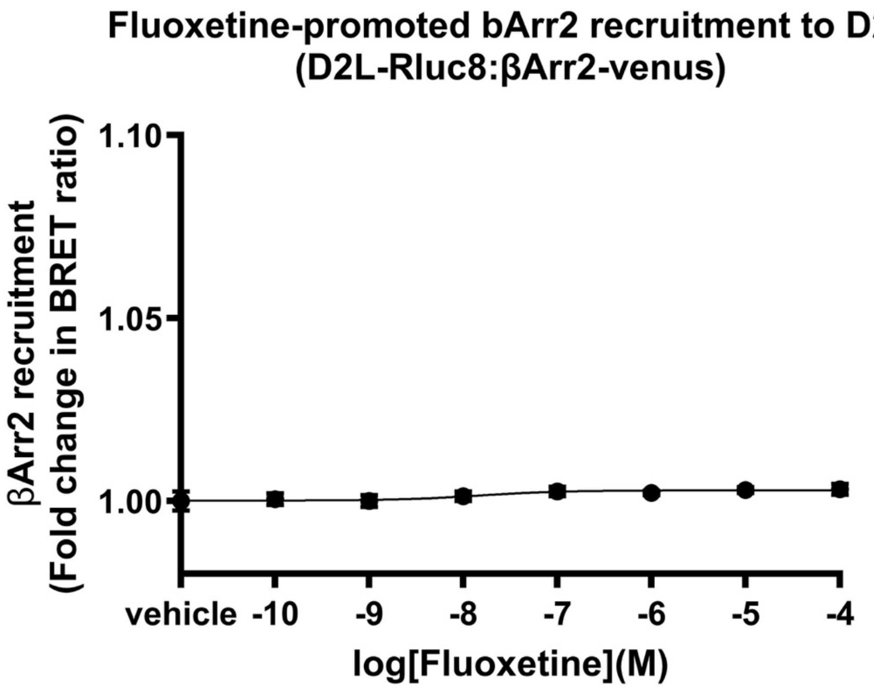
